# Decoding the interactome for Cyclic-di-AMP–producing enzyme diadenylate cyclase (DacA)

**DOI:** 10.64898/2025.12.11.693769

**Authors:** Rong Mu, Baotong Xie, Stephanie S. Momeni, Hui Wu

**Affiliations:** Department of Biomaterial and Biomedical Sciences, School of Dentistry, Oregon Health and Science University, Portland, Oregon, USA

**Keywords:** Diadenylate Cyclase, *Streptococcus mutans*, protein-protein interaction

## Abstract

Diadenylate cyclase (DacA) synthesizes the second messenger cyclic di-AMP (c-di-AMP), which regulates essential cellular processes across many Gram-positive and select Gram-negative bacteria/archaeal lineages. Although DacA is known to interact with regulators such as GlmM and CdaR, the breadth and functional relevance of its interactome remains poorly defined. Our study seeks to identify novel protein-protein interactions to further elucidate their unknown regulatory mechanisms and cellular roles. Using *Streptococcus mutans* as a model, we engineered a Flag-tagged strain (DacA-FLAG) and then performed co-immunoprecipitation under non-crosslinked and crosslinked conditions followed by mass spectrometry. We identified 22 candidates interacting proteins in non-crosslinked samples, 18 in crosslinked samples, and 6 shared between conditions. Selected partners were validated *in vivo* using split luciferase complementation. Notably, SMU_723 emerged as a key binding partner. AlphaFold-guided modeling predicated a direct DacA and SMU_723 interaction interface involving threonine 147, glutamine 148, and threonine 149 in DacA . Site-directed mutagenesis of these residues impaired binding, confirming their critical role. An SMU_723 deletion phenocopied a *dacA* deletion strain, sharing prolonged lag phase, aberrant cell morphology, reduced acid production and acid tolerance, impaired sorbitol metabolism, decreased colonization in a Drosophila model, and delayed growth upon calcium stimulation. These shared phenotypes suggest a functional and possibly regulatory link between DacA and SMU_723. Given SMU_723 sequence homology consistent with a calcium transporter, these data suggest that the DacA–723 interaction contributes to calcium homeostasis and/or calcium-responsive signaling. Collectively, our findings expand the DacA interaction network and implicate calcium transport in c-di-AMP-mediated regulation in *S. mutans*.

**Importance:** Mapping the DacA interactome reveals how environmental and intracellular cues tune c-di-AMP signaling to control stress adaptation, ion balance, and virulence traits in *S. mutans*. By identifying new DacA-associated proteins and validating SMU_723 as a previously unrecognized interactor with genetic and phenotypic linkage to DacA, this study provides new insights into the mechanistic framework for c-di-AMP regulation in *S. mutans*. The connection between DacA and a putative calcium transporter highlights a plausible axis that couples second-messenger signaling to calcium homeostasis, with implications for biofilm physiology and pathogenesis. These insights open new avenues to therapeutically modulate c-di-AMP pathways.

## Introduction

Cyclic dimeric adenosine monophosphate (c-di-AMP) (1) is a pivotal bacterial second messenger that regulates core physiological processes, including potassium homeostasis, osmotic stress responses, cell wall integrity, and virulence (2–4). Its synthesis is catalyzed by diadenylate cyclases (DACs), which convert two ATP molecules into c-di-AMP. Across bacteria and some archaea, DACs comprise multiple families distinguished by domain architecture and phylogeny. The CdaA family (hereafter) is among the most widespread and best characterized: it typically contains an N-terminal transmembrane (TM) domain (Pfam PF19293) that anchors the enzyme in the cytoplasmic membrane and a conserved C-terminal catalytic domain (Pfam PF02457) that converts ATP molecules into c-di-AMP (5).

CdaA is the sole diadenylate cyclase in streptococci, making it a non-redundant enzyme for c-di-AMP production. Genetic and physiological studies in pathogenic streptococci underscore its centrality. In *Streptococcus pneumoniae*—a leading cause of community-acquired pneumonia, CdaA is essential for viability; perturbing c-di-AMP levels alters growth and attenuates virulence (6, 7). In *Group A Streptococcus* (GAS), a highly adaptable human pathogen responsible for a wide range of illnesses—from mild pharyngitis to severe diseases such as rheumatic heart disease (RHD) and invasive infections (8)—DacA deletion produces a prolonged lag phase, abolishes biofilm formation, heightens sensitivity to osmotic, acid, and oxidative stresses, increases susceptibility to cell wall–active antibiotics, and reduces virulence (9). In *Streptococcus agalactiae*, the most common cause of culture-confirmed neonatal bacterial infections in the United States and a major global contributor to neonatal morbidity (10), DacA is required for aerobic growth and supports optimal anaerobic growth (11). In *Streptococcus mutans*, a key etiological agent of dental caries and a major contributor to acid production and biofilm formation in dental plaque (12), DacA deletion prolongs lag phase, increases autolysis, impairs oxidative stress tolerance, and disrupts biofilm formation and extracellular polymeric substance (EPS) production (13, 14). Perturbation of c-di-AMP homeostasis similarly impacts biofilm formation, adherence, stress tolerance, and virulence in *Streptococcus suis*, a zoonotic pathogen that can cause severe infections in humans (15), and in *Streptococcus gallolyticus subsp. gallolyticus* (Sgg), an opportunistic pathogen primarily associated with septicemia and infective endocarditis in the elderly and immunocompromised—and strongly linked to colorectal cancer (CRC) (16). Taken together, these findings—highlight conserved yet species-specific roles for this second messenger (17).

Although DAC’s enzymatic function is clear, how its activity is regulated in cells remains incompletely understood. Work in *Listeria monocytogenes* and *Bacillus subtilis* established that DacA engages protein–protein interactions with partners encoded in the same operon—most notably CdaR and GlmM—to tune its enzymatic activity (18, 19). In contrast, potential interactors outside the dacA operon and the broader DacA interactome in streptococci remain poorly defined. Given that c-di-AMP is essential in some streptococcal species but dispensable in others—reflecting differences in metabolism and environmental niches (11, 20, 21) — mapping the DacA interaction network is likely to reveal species-specific regulatory logic, potential crosstalk with other signaling and metabolic pathways, and new nodes for therapeutic intervention (11, 20, 21).

Here, we identified DacA-associated proteins in *S. mutans* using combined affinity purification–mass spectrometry (AP–MS) and an *in vivo* split-luciferase complementation assay. We generated a DacA-FLAG strain and identified multiple candidate interactors under both non-crosslinked and crosslinked conditions. SMU_723 was identified as a previously unrecognized DacA binding partner. Structural modeling suggested a direct DacA–SMU_723 interface mediated by residues Thr147, Gln148, and Thr149 in DacA’s catalytic domain; site-directed mutagenesis confirmed their importance for binding. Deletion of SMU_723 phenocopied multiple features of a Δ*dacA* strain such as prolonged lag phase, aberrant cell morphology, reduced acid production and acid tolerance, impaired sorbitol utilization, diminished colonization in a *Drosophila* model, and delayed growth upon calcium stimulation, pointing to a functional linkage. SMU_723 is a putative calcium transporter, suggesting that DacA may interface with calcium homeostasis or calcium-responsive signaling. Together, these findings expand the DacA regulatory landscape in *S. mutans* and implicate calcium transport as a previously underappreciated dimension of c-di-AMP control.

## Results

### Creating an affinity purification system in *S. mutans*

To identify potential protein-protein interactions involving DacA, we constructed a *dacA*-FLAG fusion strain in *Streptococcus mutans*, in which a FLAG epitope tag was fused to the C-terminus of DacA via a flexible linker to preserve structural integrity (Fig. 1A). Both non-crosslinked and crosslinked co-immunoprecipitation experiments were performed using mid-log phase cultures of the wild-type *S. mutans* UA159 strain and the engineered DacA-FLAG strain. DacA-containing protein complexes were isolated from cell lysates using anti-FLAG affinity resin, and the immunoprecipitated complexes were subject to mass spectrometry analysis. Specific enrichment of potential interactor peptides was assessed based on spectral count values, representing protein abundance. Proteins identified in the DacA-FLAG strain were compared to those from the wild-type strain lacking the FLAG tag. In total, 22 proteins from the non-crosslinked condition (Fig. 1B) and 18 proteins from the crosslinked condition (Fig. 1C) showed at least a 2-fold enrichment and had spectral counts greater than 10 in the experimental samples. Six proteins were identified in both conditions. Notably, known DacA regulators within the same genetic locus, such as GlmM and CdaR, were also detected with strong enrichment in the crosslinked and non-crosslinked samples, respectively. Adjacent to these regulators, SMU_723 emerged as a highly enriched protein that was consistently co-purified with DacA under both conditions and showed strong enrichment relative to the no-FLAG control, suggesting it is a specific interaction partner of DacA.

**Fig. 1.**
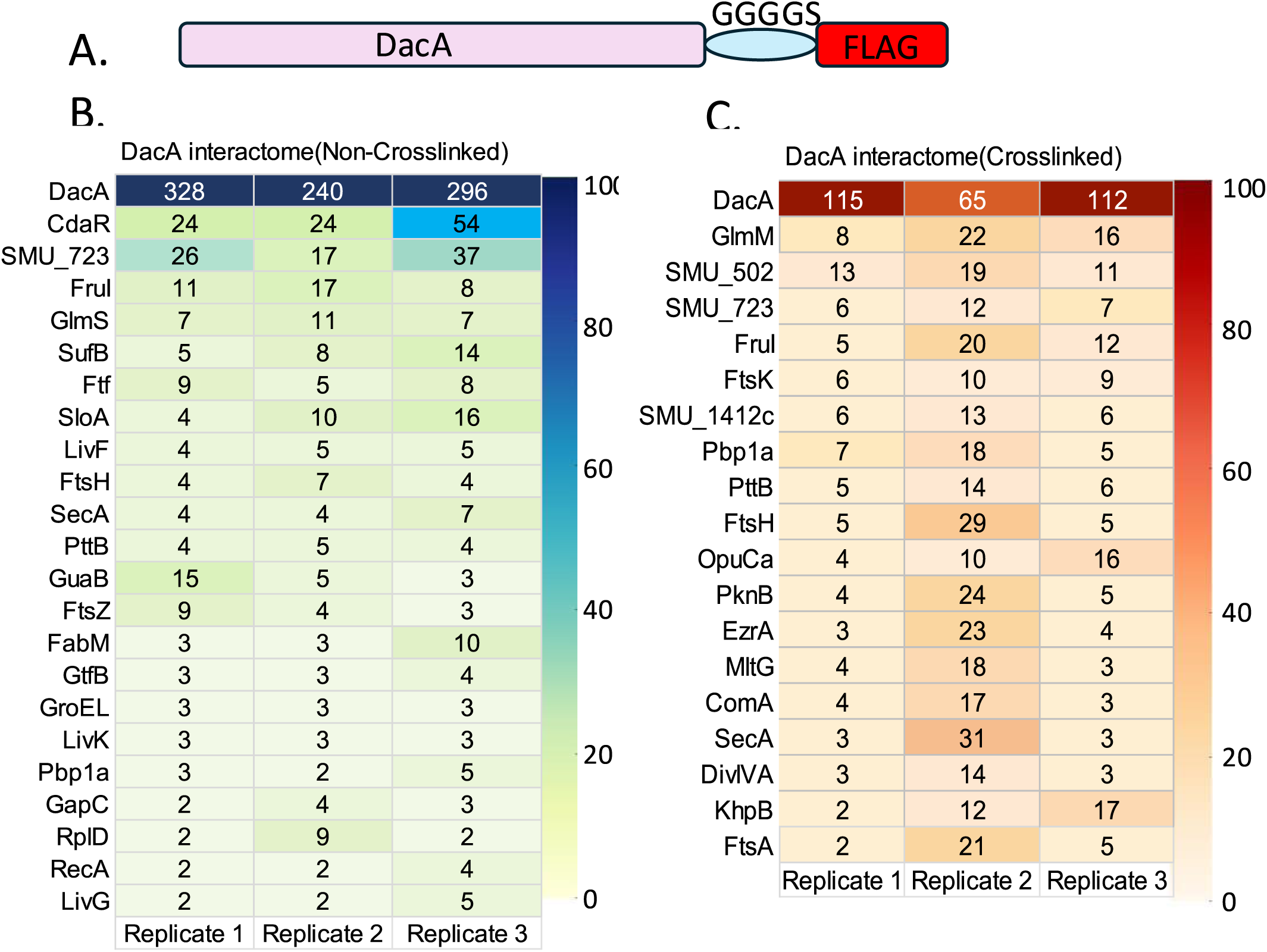
Engineering FLAG-tagged DacA and proteomic analysis of DacA interactome under non-crosslinking and crosslinking conditions. (A) Schematic representation of the FLAG-tagged DacA construct. A FLAG epitope tag was fused to the C-terminus of DacA via a flexible GGGGS linker to facilitate co-immunoprecipitation. (B) Heat map showing fold changes of proteins co-immunoprecipitated with FLAG-DacA under non-crosslinking conditions. (C) Heat map showing fold changes of proteins co-immunoprecipitated with FLAG-DacA under crosslinking conditions. Only proteins enriched ≥2-fold and with spectral counts >10 in the experimental samples were considered for analysis.

### Validation of protein-protein interaction using split luciferase complementation assay (SLCA)

Our mass spectrometry data suggested a potential moonlighting function of DacA through its interactions with various proteins of diverse functions. To validate these interactions in vivo, we employed a split luciferase complementation assay (SLCA) and selected several candidates with differing levels of enrichment from the proteomic analysis, including SMU_723, FruI, Ftf, SecA, and GapC. The N-terminal domain of Renilla luciferase (amino acids 1–155) was fused to the C-terminus of DacA, while the C-terminal domain (amino acids 156–314) was fused to the C-terminus of each candidate interactor, generating strains such as DacA-luciN+723-luciC, DacA-luciN+FruI-luciC, DacA-luciN+Ftf-luciC, DacA-luciN+SecA-luciC, and DacA-luciN+GapC-luciC. As a positive control, we constructed a DacA-luciN+GlmM-luciC fusion strain using GlmM (SMU_1426c), a known interactor of DacA. To account for non-specific interactions, we included several negative controls: (i) DacA-luciN, expressing only the N-terminal luciferase domain fused to DacA; (ii) 723-luciC, expressing only the C-terminal luciferase domain fused to SMU_723; and (iii) DacA-luciN+luciC, in which the luciferase C-terminal domain was expressed alone without a fusion partner. The SLCA results indicated that DacA specifically interacts with most of the tested candidates *in vivo*, including SMU_723, FruI, SecA, and GapC, as shown by their increased luminescence signals (Fig 2). Of note, the DacA-luciN+Ftf-luciC strain did not exhibit a significantly increased luminescence signal compared to the DacA-luciN+luciC negative control, suggesting a possible indirect or weak interaction between DacA and Ftf.

**Fig. 2.**
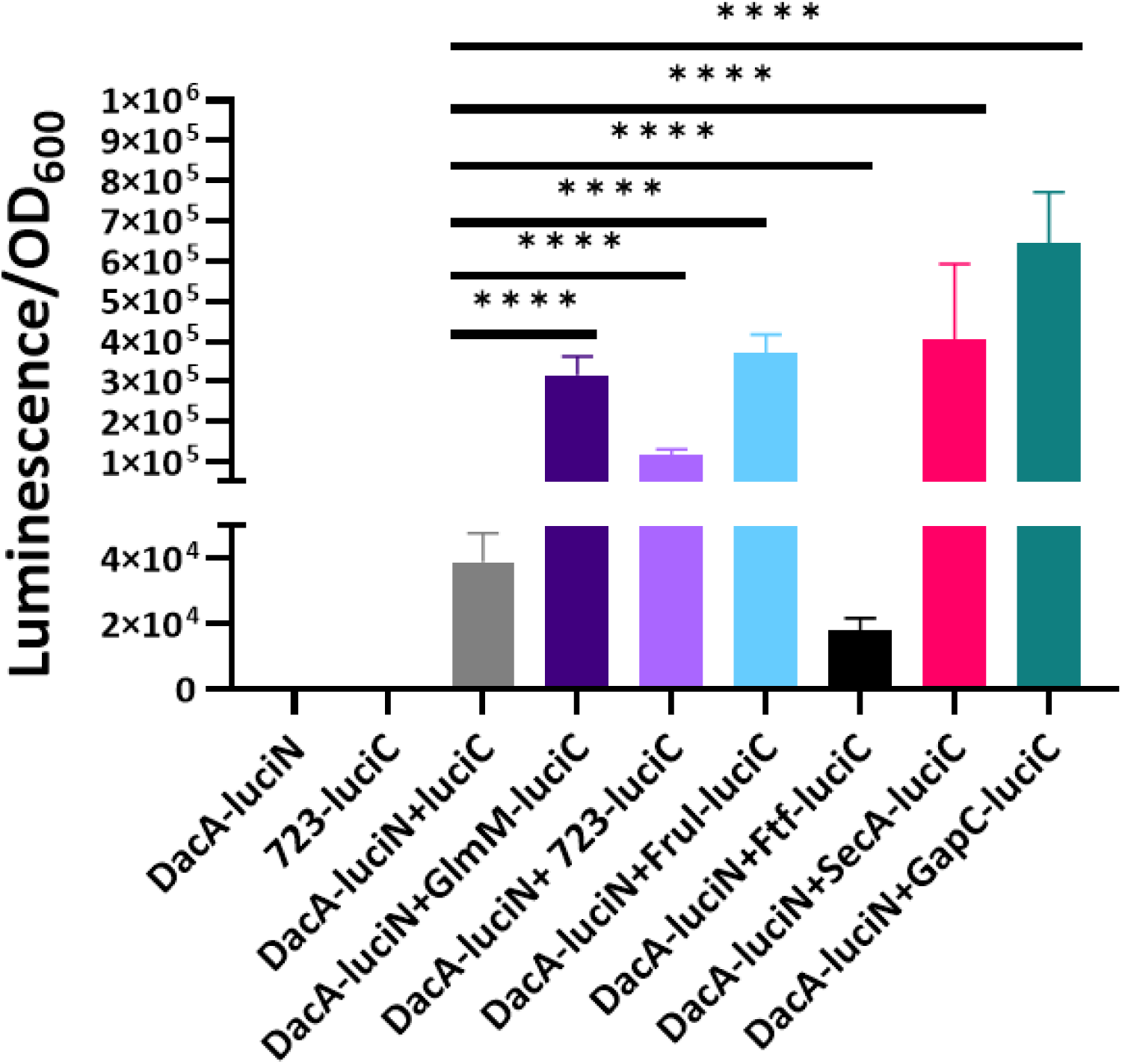
*In vivo* validation of DacA interactors using split luciferase complementation. The N-terminal domain of RenG luciferase (amino acids 1–155) was fused to the C-terminus of DacA, while the C-terminal domain (amino acids 156–314) was fused to the C-terminus of candidate DacA-interacting proteins to generate test strains: DacA-luciN+723-luciC, DacA-luciN+FruI-luciC, DacA-luciN+Ftf-luciC, DacA-luciN+SecA-luciC, DacA-luciN+GapC-luciC, and DacA-luciN+GlmM-luciC. Negative controls included: (i) DacA-luciN (N-terminal luciferase domain alone fused to DacA), (ii) 723-luciC (C-terminal domain alone fused to SMU_723), and (iii) DacA-luciN+luciC (C-terminal domain expressed alone without a fusion partner). Luminescence signals were normalized to optical density at 600 nm (OD₆₀₀). Data (three independent assays) are presented as mean ± SD and analyzed using an unpaired Student’s t-test (*P < 0.05; **P < 0.01; ***P < 0.001; **P < 0.0001).

### Assessment of cell morphology following deletion of DacA-interacting proteins

To explore the functional significance of DacA and its potential interacting partners, we selected several highly enriched candidates-SMU_502, SMU_723, FruI, and Ftf-for gene deletion analysis. We generated the corresponding mutants (Δ502, Δ723, Δ*fruI*, and Δ*ftf*) and included a previously characterized *dacA* deletion strain (Δ*dacA*) (13) for comparative phenotypic analysis. By examining these mutants, we aimed to identify shared phenotypic traits that might reflect functional connections between DacA and its interacting proteins. Growth curve analysis revealed that the ΔdacA strain exhibited a markedly prolonged lag phase, while the Δ723 mutant showed a moderately extended lag phase compared to the wild-type strain UA159 (Fig. 3A). In contrast, Δ502 and Δ*ftf* mutants displayed growth rates comparable to the wild-type strain, whereas Δ*fruI* exhibited an accelerated growth rate. These findings suggest that both DacA and SMU_723 positively contribute to growth, while FruI may be involved in a rate-limiting step in growth kinetics. Microscopic analysis further revealed distinct morphological abnormalities in the ΔdacA and Δ723 strains. When cultured in THB liquid medium, both mutants exhibited irregular cell shapes, in contrast to the uniform rod-like morphology observed in UA159 and the other deletion strains (Fig. 3B). Notably, cell morphology was restored in the complemented strains, confirming that the observed defects were specifically due to the loss of the respective genes. These observations indicate that DacA and SMU_723 play important roles in maintaining proper cell morphology.

**Fig. 3.**
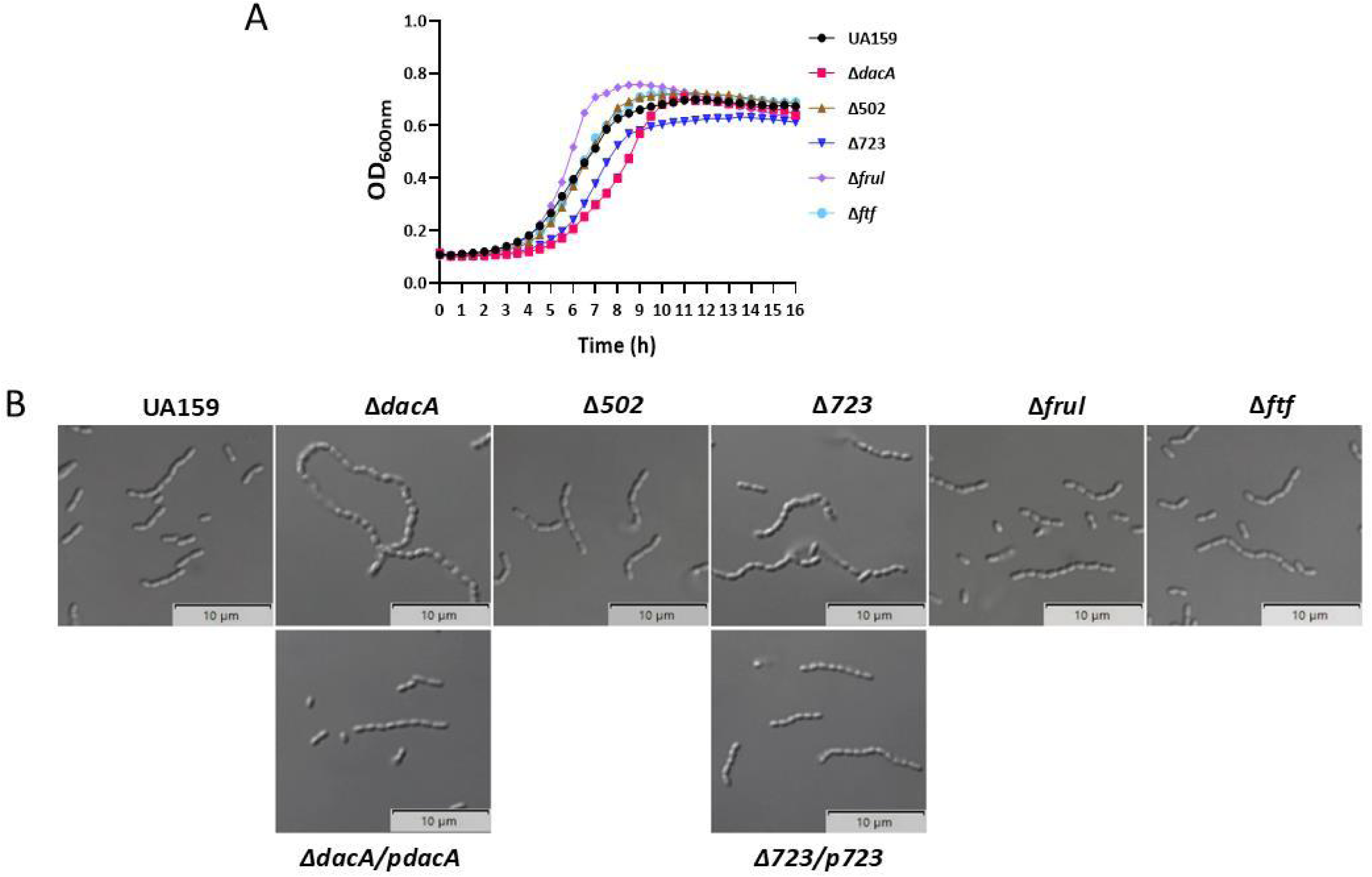
Growth and cell morphology changes in deletion strains of *S. mutans*. (A) Growth curves of wild-type strain UA159 and deletion mutants (Δ*dacA*, Δ502, Δ723, Δ*fruI* and Δ*ftf*) were measured in THB medium using a 96-well plate format. The figure presents a representative result from three independent assays. (B) Cell morphology of overnight cultures observed under 100× oil immersion microscopy. Scale bar = 10µm.

### Functional role of DacA interactors in biofilm formation

Given that biofilm formation is a key aspect of *S. mutans* pathogenicity and that the Δ*dacA* strain exhibits significant biofilm phenotype, we next investigated how deletion of DacA-interacting candidates affects this phenotype. Our results showed that only the ΔdacA strain formed biofilms with a markedly altered pattern compared to the wild-type and other mutant strains (Δ502, Δ723, Δ*fruI*, and Δ*ftf)* (Fig. 4A). Crystal violet staining further revealed that both Δ*dacA* and Δ723 mutants produced significantly less biomass relative to the wild-type strain, while no significant differences were observed in the Δ502, Δ*fruI*, and Δ*ftf* mutants (Fig. 4B). These findings suggest that DacA and SMU_723 play important roles in supporting robust biofilm formation in *S. mutans*.

**Fig. 4.**
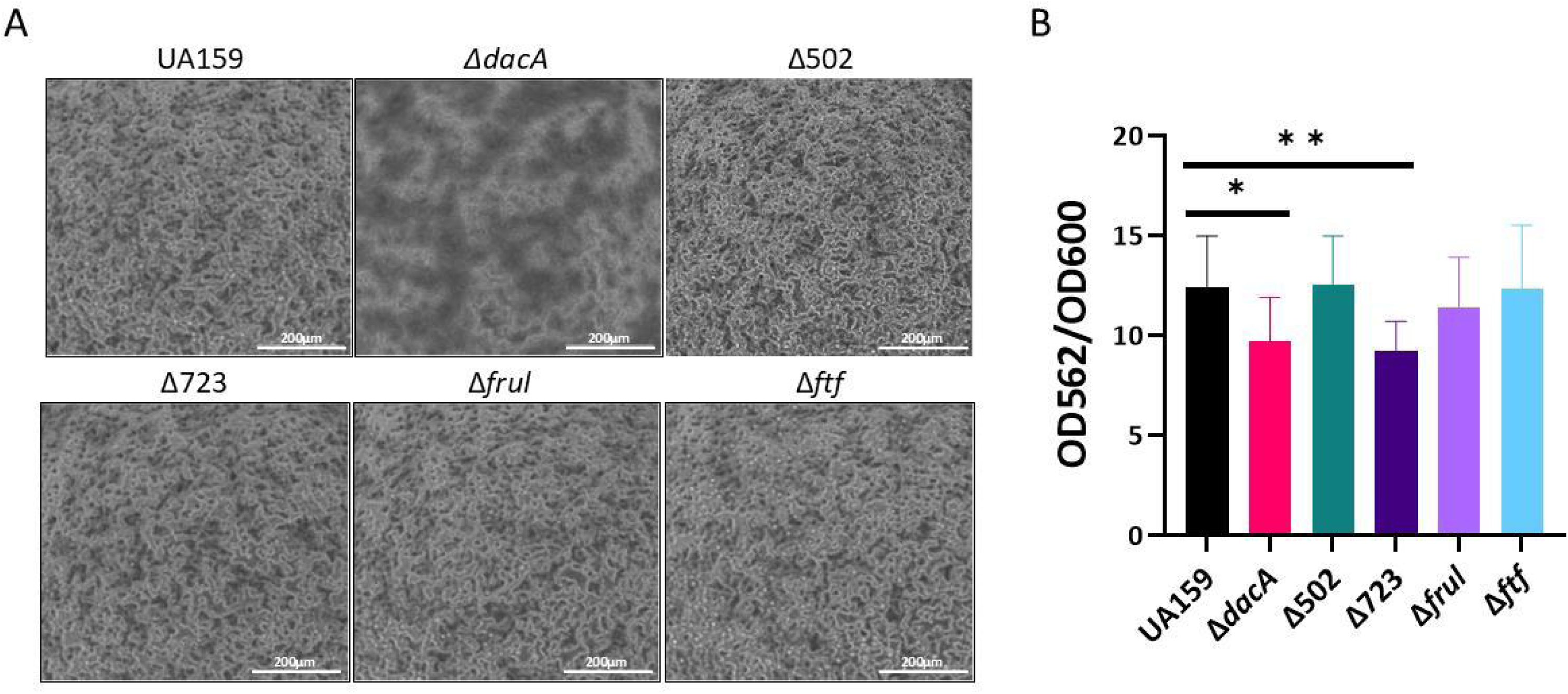
Biofilm formation and quantification. (A) Representative phase-contrast images of biofilms formed by wild-type strain UA159 and deletion mutants (Δ*dacA*, Δ502, Δ723, Δ*fruI*, and Δ*ftf*) grown in THB medium supplemented with 1% sucrose. Images were captured using a Cytation 5 Cell Imaging Multimode Reader at 40× magnification. Scale bar = 200 µm. (B) Biofilm biomass was quantified by crystal violet staining. Absorbance at 562 nm was normalized to OD₆₀₀ values. The figure presents a representative result from three independent assays. Data are presented as mean ± SD and analyzed using an unpaired Student’s t-test (*P < 0.05; **P < 0.01; ***P < 0.001; **P < 0.0001).

### Functional impact of DacA interactors on acid resistance

*S. mutans* is an aciduric bacterium capable of surviving and thriving in low-pH environments. To further investigate the role of DacA and its interacting partners in acid stress response, we examined acid tolerance in the deletion mutants. Log-phase cultures were exposed to an acid shock treatment using 0.1 M glycine-HCl buffer at pH 2.8. Following this exposure, the Δ*dacA*, Δ723, and Δ*ftf* strains exhibited significantly reduced viable cell counts compared to the wild-type strain UA159 (Fig. 5), indicating impaired survival under acidic conditions. Interestingly, the Δ502 and Δ*fruI* mutants showed significantly increased viable cell counts relative to UA159 (Fig. 5), suggesting a potential repressive role for these genes in acid stress adaptation. These findings suggest that DacA, SMU_723 and Ftf are involved in promoting *S. mutans* acid resistance. The enhanced survival of Δ502 and Δ*fruI* mutants under acid stress also raises the possibility that deletion of these genes may relieve negative regulation of acid tolerance mechanisms.

**Fig. 5.**
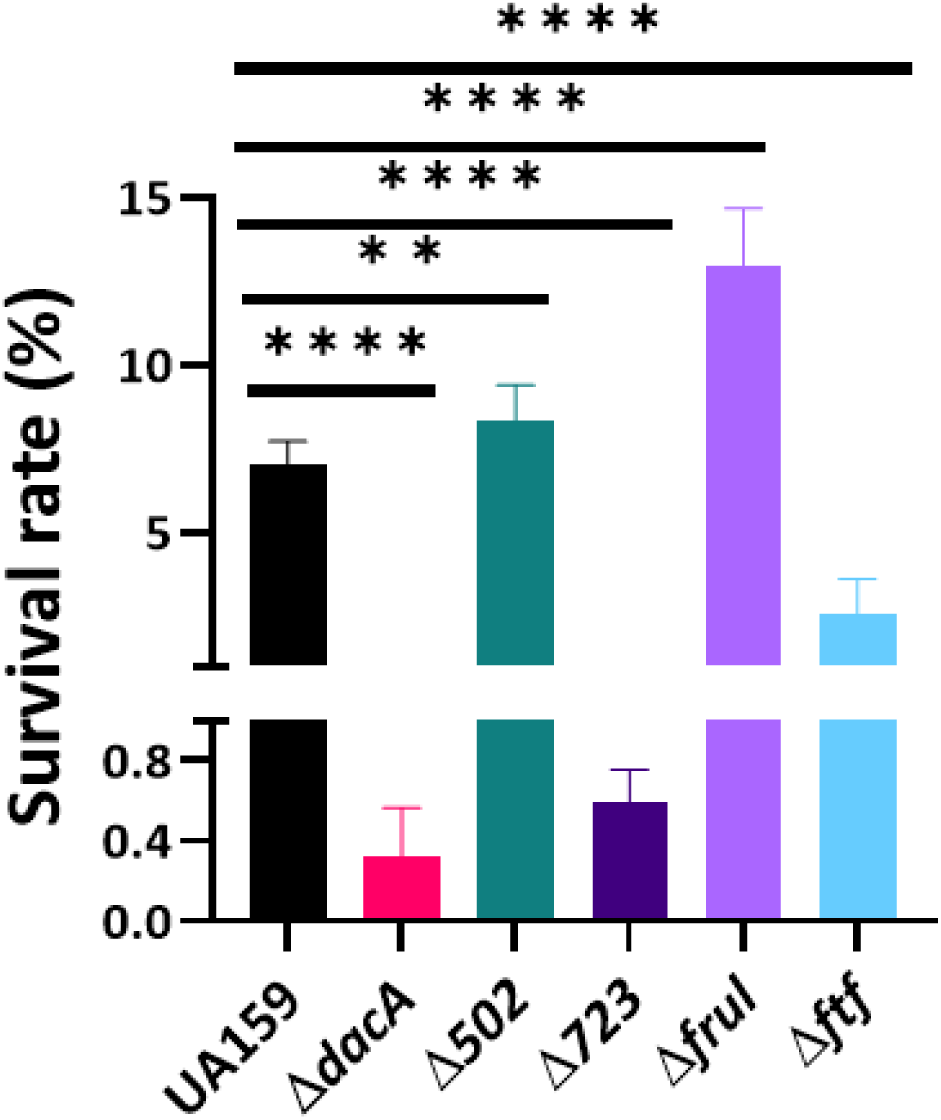
Acid shock response. Acid resistance of *S. mutans* wild-type strain UA159 and its deletion mutants (Δ*dacA*, Δ502, Δ723, Δ*fruI*, and Δ*ftf*) were assessed by exposing mid-log phase cultures to 0.1 M glycine-HCl buffer (pH 2.8) or control buffer (pH 7.0) for 5 minutes at 37°C, followed by dilution, plating on THB agar, and colony enumeration after 48 hours of incubation. Survival rates were calculated as the ratio of colony-forming units (CFUs) from the acid-treated samples to the pH 7.0 control. The figure presents a representative result from three independent assays. Data are presented as mean ± SD and analyzed using an unpaired Student’s t-test (*P < 0.05; **P < 0.01; ***P < 0.001; **P < 0.0001).

### Functional Impact of DacA interactors on sugar utilization

Given that *S. mutans* utilizes a wide range of carbohydrates to produce acid and form biofilms, we investigated whether deletion of *dacA* and SMU_723 impacts sugar metabolism. To assess this, we performed sugar utilization profiling using the API 20 Strep system. Both the Δ*dacA* and Δ723 mutants showed significantly impaired sorbitol metabolism at the 24-hour time point compared to the wild-type strain, suggesting a delayed metabolic response (Fig. 6). By contrast, the Δ502, Δ*fruI*, and Δ*ftf* mutants did not exhibit noticeable differences in sugar utilization. These findings indicate that DacA and SMU_723 are important for efficient sorbitol metabolism, which may be essential for rapid environmental adaptation and the pathogenic potential of *S. mutans* in the oral cavity.

**Fig. 6.**
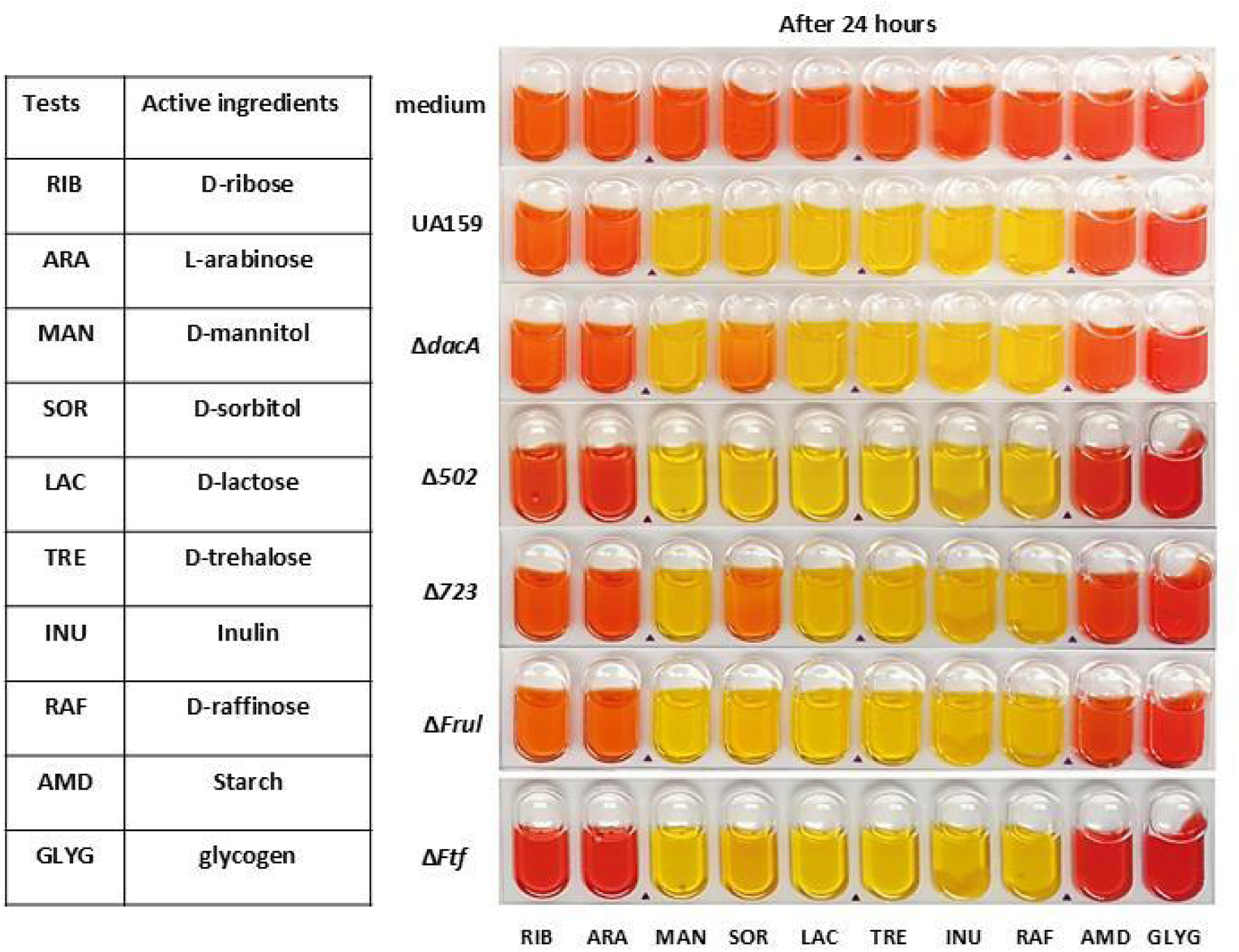
Sugar utilization profiling. Metabolic activity of *S. mutans* wild-type strain UA159 and deletion mutants (Δ*dacA*, Δ502, Δ723, Δ*fruI*, and Δ*ftf*) were assessed at 24-hour time point using the API 20 Strep system. Medium alone served as a negative control to determine which sugars were metabolized by bacterial strains. The figure presents a representative result from three independent assays.

### Deletion of *dacA* or SMU_723 reduces acid production

Given the substantial overlap in phenotypic and functional changes observed in the ΔdacA and ΔSMU_723 mutants, we sought to further investigate the biological role of SMU_723 in relation to DacAin acid production. We performed a glycolytic pH drop assay using 1% glucose and 1% sucrose as substrates. The wild-type strain UA159 exhibited the most rapid and pronounced pH decrease within the first 15 minutes. In contrast, the Δ723 mutant showed a moderate reduction in pH, while the Δ*dacA* strain displayed the least pH drop (Fig. 7A & 7B). These results suggest that both DacA and SMU_723 contribute to acid production to varying extents.

**Fig. 7.**
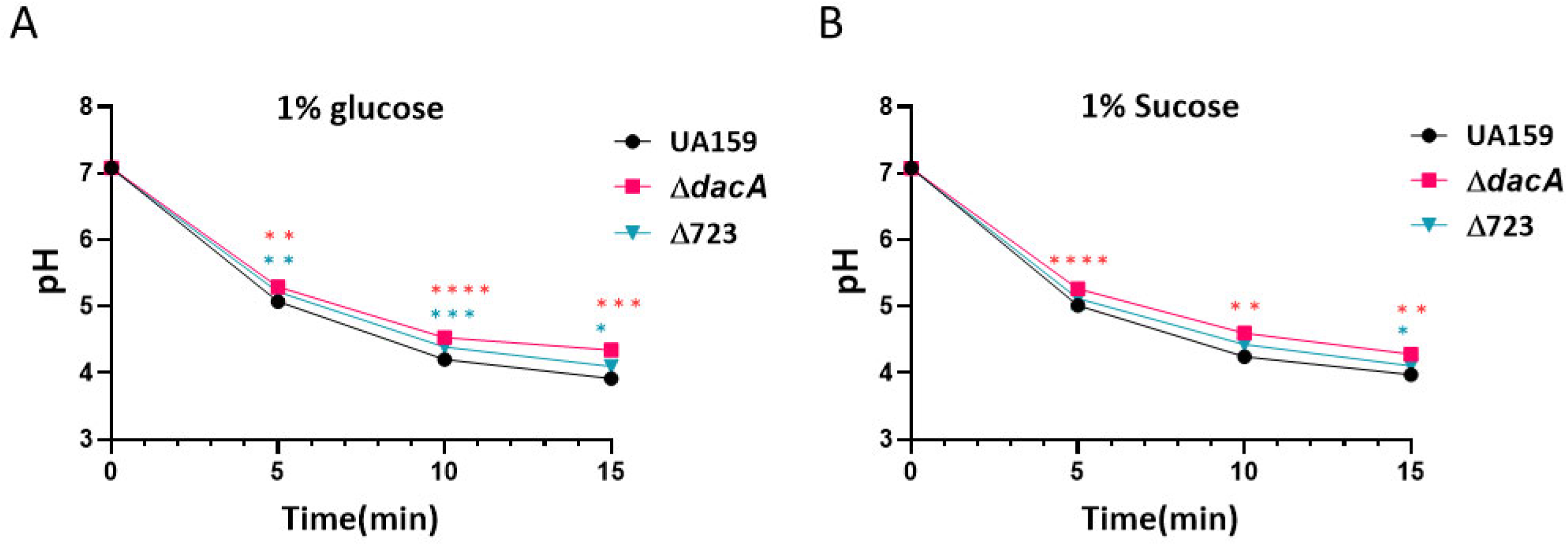
Glycolytic activity. Glycolytic pH drop assays were performed using 1% glucose (A) and 1% sucrose (B) as substrates. *S. mutans* wild-type strain UA159 and deletion mutants (ΔdacA and Δ723) were assessed. pH measurements over time were analyzed using two-way ANOVA followed by Tukey’s post hoc test. The figures present a representative result from three independent assays. Data are presented as mean ± SD (*P < 0.05; **P < 0.01; ***P < 0.001; **P < 0.0001).

### Deletion of *dacA* or SMU_723 reduces *S. mutans* colonization in *Drosophila melanogaster*

To evaluate the role of DacA and SMU_723 in bacterial colonization *in vivo*, we used *Drosophila melanogaster* as a model host. Flies from all experimental groups consumed comparable amounts of 5% sucrose, with or without bacterial supplementation, over a 3-day feeding period. Despite similar intake, colonization levels were significantly lower in flies fed with either the Δ*dacA* or Δ723 mutant strains compared to those fed with the wild-type strain (Fig. 8A & 8B). These results indicate that both DacA and SMU_723 are important for effective colonization of *S. mutans in vivo*.

**Fig. 8.**
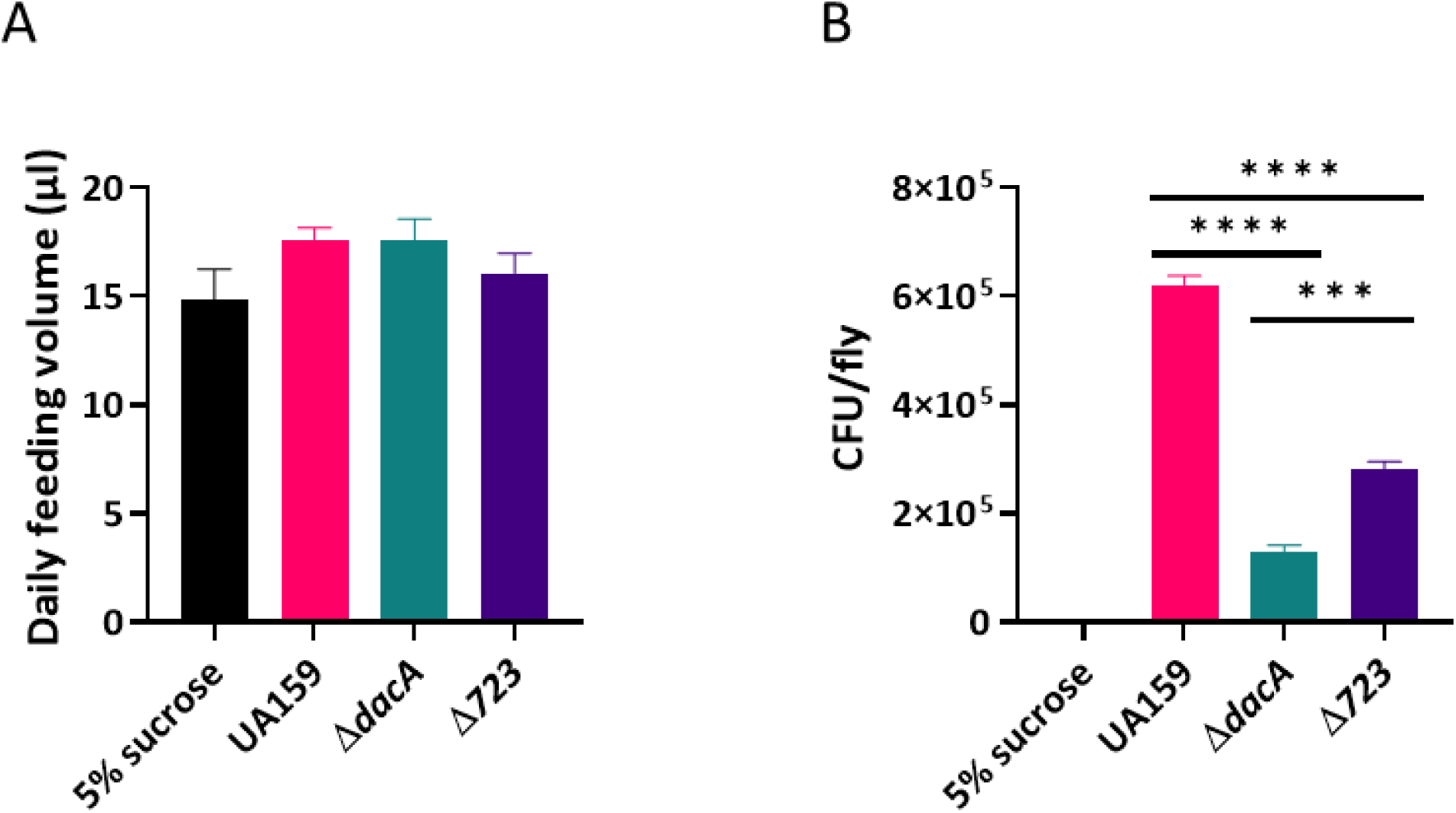
Bacterial colonization in *Drosophila in vivo*. *S. mutans* wild-type strain UA159 and its deletion mutants (Δ*dacA* and Δ723) were assessed. (A) Daily feeding volume per group was measured over three days (9 flies per group). (B) Bacterial colonization levels were determined by colony-forming unit (CFU) counts. For each group, samples were plated in triplicate to assess variation. Data are presented as mean ± SD and were analyzed using an unpaired Student’s t-test (P < 0.05; P < 0.01; P < 0.001; P < 0.0001). The figure shows one representative result from three independent experiments.

### Threonine 147, Glutamine 148, and Threonine 149 in *S. mutans* DacA are critical for interaction with SMU_723

To further validate the *in vivo* interaction between DacA and SMU_723, we used AlphaFold3 to predict the structure of their protein complex (Fig. 9A). The model suggested a potential interaction interface involving residues Threonine 147, Glutamine 148, and Threonine 149 in DacA. To assess the functional importance of these residues, we generated a DacA alanine substitution variant (DacA3A), in which T147, Q148, and T149 were replaced with alanine. This mutant was fused to the luciferase N-terminal domain (amino acids 1–155) and paired with the luciferase C-terminal domain (amino acids 156–314) fused to the C-terminus of SMU_723, generating the DacA3A-luciN+723-luciC strain. Luminescence from the DacA3A-luciN+723-luciC strain was significantly reduced compared to the DacA-luciN+723-luciC strain (Fig. 9B), indicating that substitution of these residues disrupts the interaction. These results demonstrate that Threonine 147, Glutamine 148, and Threonine 149 are critical for DacA–SMU_723 binding and further support the existence of a direct *in vivo* interaction between the two proteins.

**Fig. 9.**
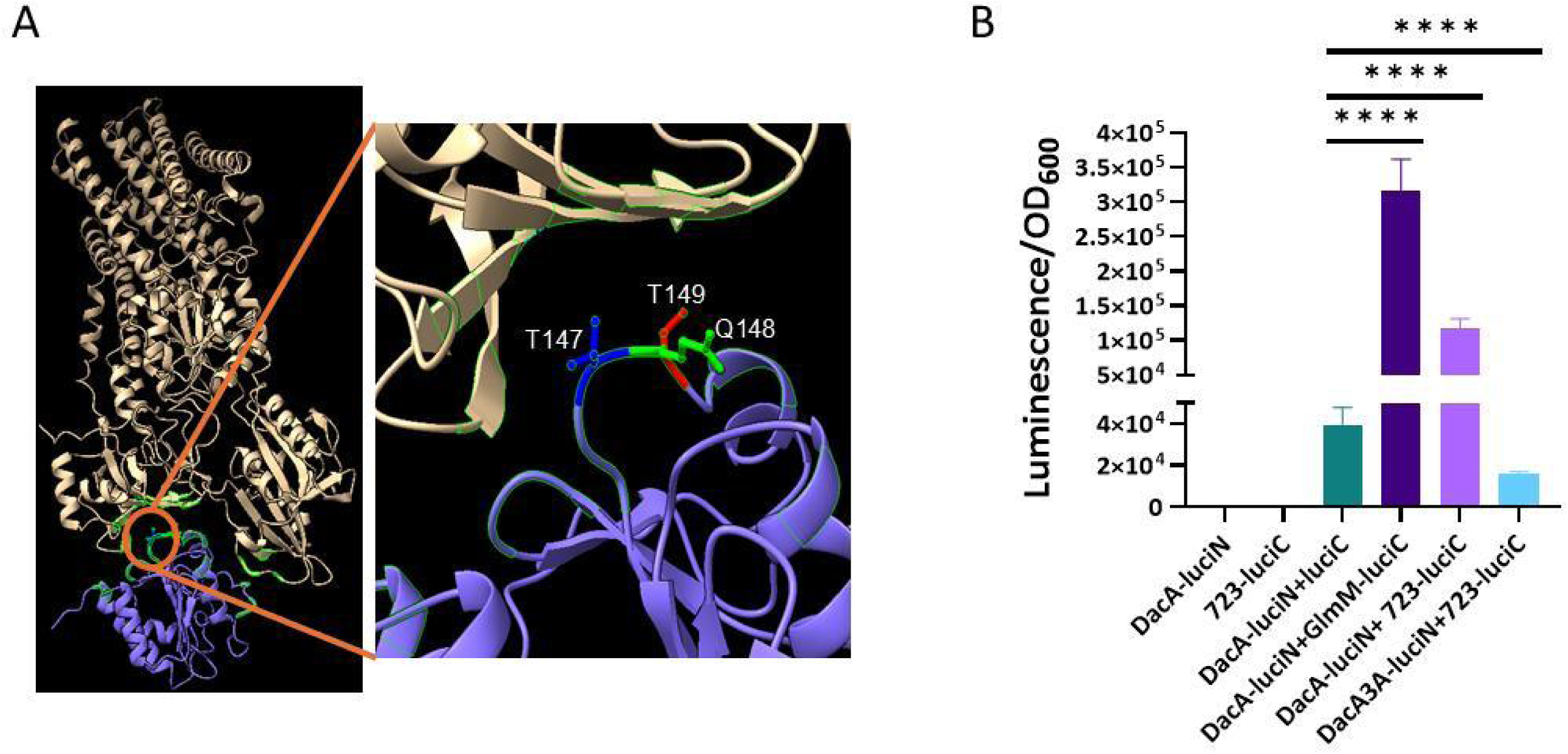
Prediction and validation of critical residues mediating the DacA–SMU_723 protein interaction. (A) The interaction between DacA (excluding its transmembrane domain) and full-length SMU_723 was predicted using AlphaFold3 (35). The resulting structural model was visualized with UCSF ChimeraX version 1.9 (36). Three residues in DacA—Threonine 147 (T147), Glutamine 148 (Q148), and Threonine 149 (T149)—were identified at the predicted interface and are highlighted. (B) To assess the functional importance of these residues, a site-directed mutant DacA3A-luciN + 723-luciC was generated by substituting T147, Q148, and T149 with alanine. This mutant was tested alongside the wild-type interaction pair DacA-luciN + 723-luciC, a positive control DacA-luciN + GlmM-luciC, and three negative controls: (i) DacA-luciN, (ii) 723-luciC, and (iii) DacA-luciN co-expressed with unfused luciC. Luminescence signals were normalized to OD₆₀₀. The figure shows a representative result from three independent experiments. Data are presented as mean ± SD and analyzed using an unpaired Student’s t-test (*P < 0.05; **P < 0.01; ***P < 0.001; ****P < 0.0001).

### DacA–SMU_723 interaction modulates calcium-dependent growth in *S. mutans*

SMU_723 encodes a predicted transmembrane protein homologous to the calcium ATPase LMCA1 from *Listeria monocytogenes*, sharing 65% amino acid identity. Recent structural studies have demonstrated that LMCA1 plays an essential role in calcium transport in *L. monocytogenes* (22). Using AlphaFold3, we predicted the structure of SMU_723 in complex with Ca²⁺ and ATP, revealing a conformation similar to that of LMCA1 (Fig. 10A). Given the direct interaction between DacA and SMU_723 identified in our assays, we hypothesized that calcium signaling may be an important component of the DacA regulatory network. To investigate this, we assessed bacterial growth in response to CaCl₂ supplementation. The wild-type *S. mutans* UA159 strain exhibited enhanced growth in the presence of 1mM calcium (Fig. 10B). In contrast, the Δ*dacA* mutant showed no enhanced growth upon calcium supplementation (Fig. 10C), suggesting a loss of calcium responsiveness. The ΔSMU_723 mutant displayed a prolonged lag phase and a slightly reduced growth rate in the presence of calcium (Fig. 10D), indicating that SMU_723 is also required for calcium-dependent growth enhancement. To further explore the functional significance of the DacA–SMU_723 interaction, we tested calcium effects on strains expressing fusion proteins used in our SLCA assay: one strain (DacA-luciN+723-luciC) contains intact DacA and SMU_723 fusion proteins, while the other (DacA3A-luciN+723-luciC) harbors a site mutation in DacA that disrupts its interaction interface with SMU_723. The DacA-luciN+723-luciC strain exhibited enhanced growth in the presence of calcium, similar to the wild-type (Fig. 10E). In contrast, the DacA3A-luciN+723-luciC strain showed a prolonged lag phase and a slightly reduced growth rate upon calcium supplementation, closely resembling the ΔSMU_723 phenotype (Fig. 10F). These findings suggest that disruption of the DacA–SMU_723 interaction weakens calcium responsiveness.

**Fig. 10.**
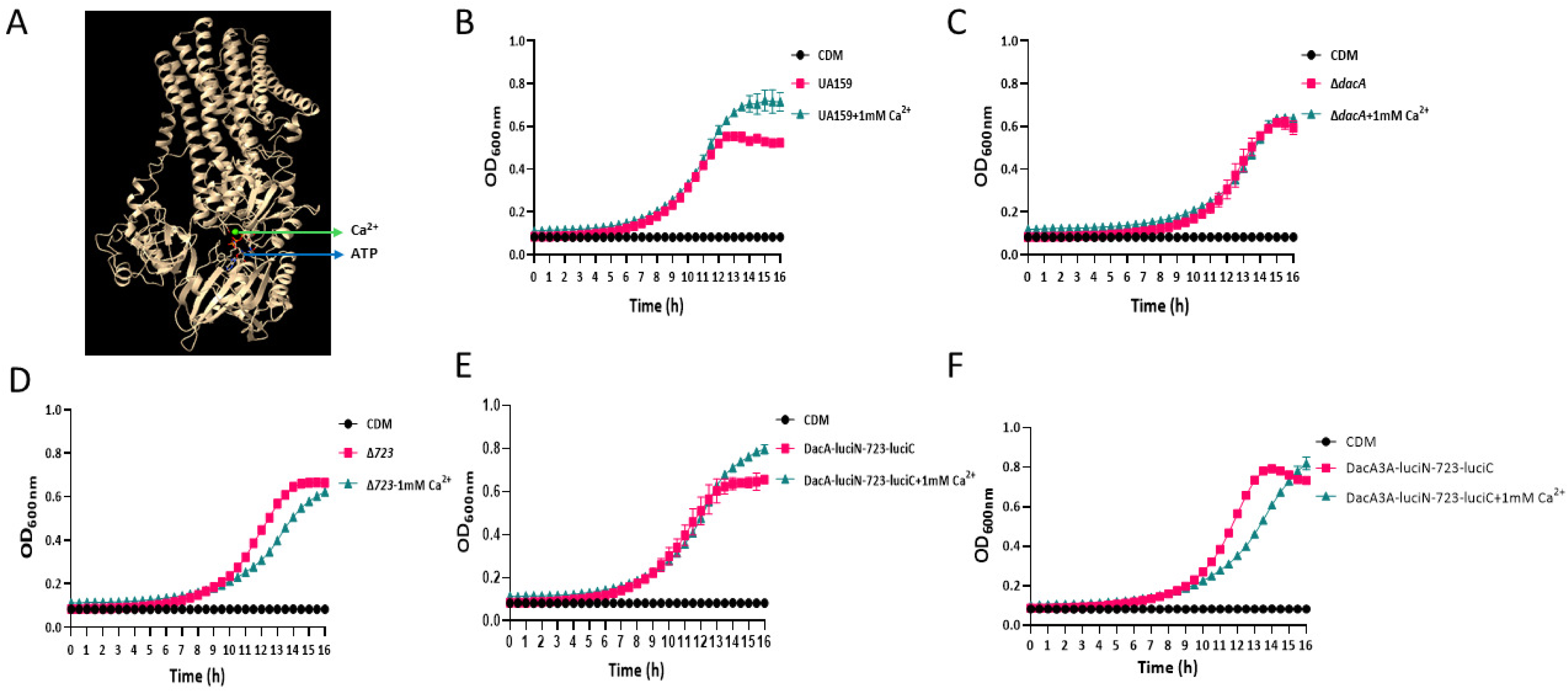
Structural prediction of SMU_723 as a calcium transporter and assessment of calcium involvement in bacterial growth regulation. (A) The full-length SMU_723 protein structure in complex with Ca²⁺ and ATP was predicted using AlphaFold3 (34). Growth curves of wild-type strain UA159 (B), deletion strain ΔdacA (C), Δ723 strain (D), DacA-luciN+723-luciC strain (E) and site mutated strain DacA3A-luciN+723-luciC (F) were measured in chemical defined media with or without 1 mM calcium. Bacteria were cultured in a 96-well plate, and the plate was incubated in the Cytation 5 Cell Imaging Multimode Reader at 37°C with 5% CO₂. Optical density at 600 nm (OD₆₀₀) was recorded over time. The figure presents a representative result from three independent assays.

## Discussion

By combining affinity purification under both non-crosslinked and crosslinked conditions with mass spectrometry-based proteomics, we captured a broad spectrum of stable and transient DacA interactions. Robust enrichment of the known partners CdaR and GlmM in our dataset internally validates our experimental approach and reaffirms conserved, operon-linked regulation of DacA associated activity. Building on this foundation, we delineated a more complex DacA protein-protein interaction network. We identified constitutive partners present in both conditions-SMU_723, FruI, FtsH, SecA, PttB and Pbp1a. Additionally, we also identified interactions preferentially detected after crosslinking-SMU_502, FtsK, SMU_1412c, OpuCa, PknB, EzrA, MltG, ComA, DivIVA, KhpB and FtsA. The differential recovery across conditions underscores the value of orthogonal capture strategies to reveal the full spectrum of protein interactions.

Functionally, the network is strikingly diverse. Identification of enzymes of central metabolism, including GlmS, GuaB, GapC, GlmM, Ftf, and GtfB in the network suggests that DacA sits at the intersection of second messenger signaling and metabolic flux. The prominence of transport-related proteins, multiple ABC transporter components (e.g., SMU_723, SloA, LivF, LivG, SMU_1412c, OpuCa, ComA) and PTS elements (e.g., FruI, PttB) is consistent with c-di-AMP’s established role in controlling ion/osmolyte homeostasis and turgor (11, 23). Identification of cell division and morphogenesis factors like FtsZ, FtsA, EzrA, DivIVA, PknB and Pbp1a implicates DacA in coordinating envelope growth with cell-cycle progression. Consistent with this, c-di-AMP functions as a master regulator of cell volume and turgor, coupling osmotic homeostasis with cell wall integrity (11, 24). Interactions with protein quality control/trafficking machinery (GroEL, FtsH, and SecA) raise the possibility that DacA activity or turnover is tuned by proteostasis. Finally, interactions with nucleic acid-associated proteins (e.g., RecA, FtsK, KhpB) hint at c-di-AMP-linked responses to DNA damage, repair and RNA metabolism. Together, these observations extend DacA’s influence well beyond c-di-AMP synthesis per se, positioning it as a signaling hub that integrates transport, envelope homeostasis, and growth. A particularly notable finding is the DacA interaction with OpuCa, an osmolyte transporter component whose orthologs are established c-di-AMP receptors in multiple bacterial species, including *S. agalactiae* (11), *Listeria monocytogenes, Enterococcus faecalis* (25), *Staphylococcus aureus* (26) and *Bacillus subtilis* (27). This dual connectivity-protein-protein association with DacA and ligand recognition of c-di-AMP supports a feedback loop or co-regulation mechanism in which c-di-AMP both (i) modulates transporter activity and (ii) is, in turn, regulated by physical proximity or scaffolding with transport machinery via tuned DacA activity. Such reciprocal coupling could stabilize intracellular osmotic balance by matching second messenger production to the status of osmolyte flux, a regulatory motif that has been proposed in other Gram-positive systems (4, 27).

Because of its strong, reproducible enrichment and conservation, we focused on SMU_723, a predicted P-type ATPase. It shows remarkable conservation across *Streptococci*, maintaining 72-81% amino acid sequence identity. Notably, it also shares significant homology to LMCA1 in *L. monocytogenes* (65% identity), a known calcium exporter critical for bacterial survival under calcium stress, supporting its role in calcium efflux. Multiple lines of evidence link SMU_723 functionally to DacA: 1) Δ723 phenocopies ΔdacA (prolonged lag phase, morphology defects, reduced acid production/resistance, impaired sorbitol metabolism, diminished *Drosophila* colonization). 2) both mutants failed to mount calcium-stimulated growth responses; 3)alanine substitutions at Thr147, Gln148, Thr149 within DacA’s catalytic domain—residues predicted by AlphaFold to contact SMU_723—attenuate binding and recapitulate delayed growth onset upon Ca²⁺ supplementation. These convergent phenotypes argue that the DacA–SMU_723 interaction promotes timely adaptation to environmental cues, with calcium emerging as a key input. From these results, we propose three, non-exclusive regulatory scenarios: (1) DacA to Smu_723 control: DacA regulates SMU_723 activity or localization, coupling c-di-AMP production to calcium export activity and envelope physiology; (2) SMU_723-dependent control of DacA: Smu_723 modulates DacA (e.g., by scaffolding, lipid-microdomain positioning, or conformational control) to adjust c-di-AMP output in response to calcium activities; or (3) Integrated hub: DacA, Smu_723, and OpuC/other transporters assemble into a novel signaling complex that coordinates c-di-AMP and calcium signaling pathways to balance turgor, cell-wall biogenesis, and growth. Future studies in mechanistic dissection of the DacA-Smu_723 interface should clarify the working model. In conclusion, our findings position DacA as a central node in *S. mutans* that coordinates second messenger signaling with ion transport, metabolism, and cell-cycle processes. The DacA–SMU_723 axis, together with OpuCa and additional transport/division partners, suggests a membrane-embedded regulatory network that couples c-di-AMP dynamics to calcium and osmotic homeostasis. The fact that both a dacA and SMU_723 deletion led to a delayed sorbitol metabolism is a strong indication that they are part of the same pathway. The observed delay suggests that without one or both components, the bacterium loses its ability to rapidly "turn on" or fully activate its sorbitol metabolic machinery. Investigating how these interactions shape virulence traits (biofilm formation, acid tolerance, bacterial colonization) may uncover new targets for selectively disrupting cariogenic physiology while minimizing impacts on commensals.

## Materials and Methods

### Bacterial strains and culture conditions

The bacterial strains and plasmids used in this study are listed in Table S1 and *S. mutans* UA159 served as the wild-type parental strain. All *S. mutans* strains were cultured in Todd Hewitt medium (Difco) or chemical defined medium (28) and incubated at 37°C in a 5% CO₂ environment. For the selection of antibiotic-resistant colonies, THB plates were supplemented with 1 mg/mL spectinomycin, 800 µg/mL kanamycin, and 12.5 µg/mL erythromycin.

### DNA manipulation and strain construction

Primers used in this study are listed in Table S2. Specific details of strain construction are described in Supplemental Experimental Procedures. Individual polymerase chain reactions (PCRs) were performed using KOD Hot Start DNA Polymerase (millipore sigma), while overlap extension PCR (OE-PCR) was performed using KOD Xtreme Hot Start DNA Polymerase (millipore sigma).

### Coimmunoprecipitation

Coimmunoprecipitation assay was performed as previously described (29). Briefly, *S. mutans* strains UA159 and DAC-FLAG were cultured at 37°C in a CO₂ incubator to mid-log phase in THB medium, with 12.5 µg/mL erythromycin added for DacA-FLAG. Cells were pelleted, washed with ice-cold phosphate-buffered saline (PBS), and divided into two groups: non-crosslinked and crosslinked samples. For crosslinked samples, cells were incubated in PBS containing 1% (vol/vol) formaldehyde for 20 min at room temperature, followed by quenching with 0.125 M glycine for 5 min. After washing and pelleting, cells were sonicated in lysis buffer (150 mM NaCl, 10% (vol/vol) glycerol, 100 mM Na-phosphate buffer (pH 7.0) and 1% Triton X-100, protease inhibitor cocktail, pH 7.0), and the supernatant obtained after cell lysis were incubated with anti-FLAG M2 affinity gel (Sigma-Aldrich) for 3 h at 4°C with end-over-end rotation. FLAG Peptide (Sigma) with a working concentration of 150 ng/µL was used for the competitive elution of FLAG tagged DacA. The wild-type strain UA159 served as a no FLAG epitope control sample.

### In-Gel Digestion and LC-MS/MS Analysis

Proteins were separated on NuPAGE 10% bis-tris gels (Invitrogen) at 150 V for 5 min and stained with Coomassie Blue. Visible bands were excised, cut into ∼1 mm² pieces, and subjected to in-gel digestion using Trypsin Gold (Promega) and ProteaseMAX surfactant (Promega) following the manufacturer’s protocol. Mass spectrometry analysis was conducted at the Oregon Health & Science University (OHSU) Proteomics Shared Resource.

### Split luciferase complementation assays

Split luciferase complementation assays were performed as previously described (30). Overnight *S. mutans* cultures were diluted 1:20 in THB and grown to mid-log phase. Luciferase activity was measured after adding 1 µL of 0.75 mg/mL coelenterazine-h solution (Fisher) to 100 µL of cell culture using a GloMAX Discover 96-well luminometer (Promega). To normalize the data, luciferase values were divided by their corresponding optical density (OD_600_) values. The experiment was performed in triplicate and repeated three times.

### Growth curve measurement

Overnight cultures of *S. mutans* strains were diluted 1:20 in fresh THB and grown to an OD_600_ of 0.4. The cultures were then further diluted 1:100 in fresh THB and incubated in a 96-well plate at 37°C with CO₂ using a Cytation 5 Cell Imaging Multimode Reader. Cell growth was monitored at OD_600_ every 30 minutes, with a 10-second shaking step at 205 cpm before each reading to ensure proper mixing. To investigate the effects of calcium on cell growth, chemically defined medium was used in place of THB, and CaCl₂ was added to a final concentration of 1 mM. The experiment was performed in triplicate and repeated three times.

### Cell morphology examination

Cell morphologies of *S. mutans* strains were examined and imaged using an Olympus IX73 inverted microscope equipped with a 100 x oil immersion objective.

### Biofilm imaging and quantification

*S. mutans* strains were cultured overnight in THB medium, subcultured into fresh THB to mid-log phase. Cells were washed with 1× PBS and adjusted to an OD₆₀₀ of 0.4 in 1× PBS. Cultures were then diluted 1:1000 into THB supplemented with 1% sucrose and inoculated into a 96-well plate. The plate was incubated for 16 hours at 37°C in a 5% CO₂ incubator. Following incubation, OD₆₀₀ was measured. Then the plate was gently washed three times with 1× PBS. Biofilm morphology was then imaged using the Cytation 5 Cell Imaging Multimode Reader at 40× phase contrast. After that, biofilms were stained with 0.1% crystal violet. After washing with 1× PBS, the bound dye was solubilized with 30% acetic acid. Absorbance was measured at 562 nm and normalized to OD₆₀₀ values. The experiment was performed in triplicate and independently repeated three times.

### Glycolytic pH drop assay

The glycolytic pH drop assay was performed as previously described (31). *S. mutans* strains were grown overnight in THB medium and subcultured until reaching an OD_600_ of 0.6. Cells were harvested, washed, and resuspended in a solution containing 50 mM KCl and 1 mM MgCl₂, adjusted to pH 7.0. Glucose and sucrose were added to the cell suspension at a final concentration of 1%, and the pH was measured every 5 minutes for 15 minutes at room temperature using a Fisherbrand accumet AB15 Basic pH meter. The experiment was performed in triplicate and repeated three times.

### Acid shock assay

*S. mutans* acid tolerance was evaluated as previously described (31). *S. mutans* strains were cultured overnight in THB medium and subsequently subcultured until reaching an OD₆₀₀ of 0.6. Cells were then pelleted, washed with 1× PBS, and resuspended in 0.1 M glycine-HCl buffer at either pH 2.8 (acid treatment) or pH 7.0 (control). The suspensions were incubated at 37°C for 5 minutes, followed by dilution in 1× PBS and plating on THB agar. After incubation at 37°C for 48 hours, colonies were counted, and the survival rate was calculated by dividing the colony-forming units (CFUs) from the acid-treated group by those from the pH 7.0 control. The experiment was performed in triplicate and repeated three times.

### Sugar metabolism test

The API 20 Strep system was used to assess the metabolic profiling of sugar utilization in *S. mutans* strains. Briefly, overnight cultures of *S. mutans* strains were subcultured in THB medium until reaching mid-log phase. Cells were then pelleted, washed with 1× PBS, and resuspended in the API 20 Strep medium to an OD_600_ of 0.4. The medium alone served as a negative control. The strips were incubated at 37°C in a CO₂ incubator, and metabolic activity was recorded after 24 hours. The experiment was repeated three times.

### *In vivo* colonization assay

*Drosophila* Canton-S was used as an *in vivo* model to assess bacterial colonization, following a modified version of previously described methods (32–34) . Briefly, healthy 3-day-old male flies were pretreated with antibiotics for two days, then grouped into vials (9 flies per vial). Flies were fed twice daily for three days with either 5% sucrose (control) or 5% sucrose containing S. mutans strains, delivered via capillaries (Fisher #NC127687). At the end of the feeding period, flies were homogenized in 200 µL of 1× PBS using a plastic pestle. Serial dilutions were plated on THB agar and incubated at 37°C for 48 hours for colony enumeration. The assay was conducted under a 12-hour light/dark cycle at 25°C in a Drosophila incubator and repeated three times.

## Data analysis

Statistical analyses were performed using Prism 10 (GraphPad Software Inc.). An unpaired two-tailed Student’s T-test was used for comparisons between two groups. For comparisons involving multiple time points, two-way ANOVA followed by Tukey’s post hoc test was applied. A P-value < 0.05 was considered statistically significant. Data are presented as mean ± SD.

## Acknowledgements

This project was supported by NIH grants DE028329 to HW.

We gratefully acknowledge the OHSU Proteomics Shared Resource for mass spectrometry analysis. We also thank Dr. Hua Qin from Dr. Justin Merritt’s laboratory for generously providing cell cultures used as DNA templates in generating constructs for split luciferase complementation assays. The authors declare no conflict of interest.

## Data availability

The UA159 *Streptococcus mutans* reference genome is publicly available in NCBI (RefSeq accession NC_004350.2). Summarized mass spectrometry data are presented in Figures 1B & 1C. All other data supporting the findings of this study are available from the corresponding author upon request.

